# Development of a highly sensitive bioanalytical assay for the quantification of favipiravir

**DOI:** 10.1101/2021.02.03.429628

**Authors:** Paul Curley, Megan Neary, Usman Arshad, Lee Tatham, Henry Pertinez, Helen Box, Rajith KR Rajoli, Anthony Valentijn, Joanne Sharp, Steve P Rannard, Andrew Owen

## Abstract

Favipiravir (FAV; T-705) has been approved for use as an anti-influenza therapeutic and has reports against a wide range of viruses (e.g., Ebola virus, rabies and norovirus). Most recently FAV has been reported to demonstrate activity against SARS-CoV-2. Repurposing opportunities have been intensively studied with only limited success to date. If successful, repurposing will allow interventions to become more rapidly available than development of new chemical entities. Pre-clinical and clinical investigations of FAV require robust, reproducible and sensitive bioanalytical assay. Here, a liquid chromatography tandem mass spectrometry assay is presented which was linear from 0.78-200 ng/mL Accuracy and precision ranged between 89% and 110%, 101% and 106%, respectively. The presented assay here has applications in both pre-clinical and clinical research and may be used to facilitate further investigations into the application of FAV against SARS-CoV-2.

## Introduction

Favipiravir (FAV; T-705) has been reported to exert broad spectrum activity against RNA viruses, and is approved for use in Japan as an oral anti-influenza treatment [1]. Within patients receiving FAV as an influenza treatment, a high genetic barrier of resistance has been observed but deployment is complicated by concerns about teratogenicity. [2] FAV acts through lethal viral mutagenesis[3] and has demonstrated efficacy within animal models against a wide number of viruses including pathogenic avian influenza H5N1 and H7N9, Lasa virus, haemorrhagic fever viruses and Ebola virus amongst others [1, 4–10]. FAV has also shown efficacy in humans in the treatment of Ebola virus, influenza, rabies, norovirus and severe fever with thrombocytopenia syndrome. [1] Most recently, favipiravir has also been demonstrated to exert anti-SARS-CoV-2 activity *in vitro* and in the Syrian Golden Hamster model of infection [11]. High intraperitoneal doses (1000 mg/kg) were required in this model but plasma C_min_ concentration at this dose were comparable to those seen in humans. Previous work by the investigators indicate that FAV plasma concentrations in humans exceed the SARS-CoV-2 *in vitro* EC_90_ (C_max_ / EC_90_ ratio) [12], but plasma concentrations of FAV diminish with time after multiple dosing [13] and antiviral concentrations identified in Vero E6 cells (EC50 ~62μM corresponding to an EC_90_ of ~159μM; [14]) are not consistently maintained across the dosing interval. Notwithstanding, recent modelling has hinted that achievable doses should maintain intracellular concentrations of the active metabolite (favipiravir ribofuranosyl-5’-triphosphate) between doses [15]. Assessment of FAV anti-SARS-CoV-2 activity in more representative cell lines (e.g. human primary or transformed lung epithelial models) will provide more confidence in the target values. However, as of 25^th^ January 2021, 53 trials were listed on clinicaltrials.gov investigating the use of FAV for COVID19 applications [16].

Currently there is a paucity of published pharmacokinetic data for FAV and very few validated LC-MS/MS methods in relevant biological matrices. There are a number of HPLC based assays that while fully validated in plasma, are unlikely to produce the sensitivity required to fully assess the plasma and tissue pharmacokinetics of FAV as a SARS-CoV-2 therapeutic [17–20]. While LC-MS/MS methods have been published they are lacking in validation in relevant biological matrices. Previous studies have examined FAV concentrations in an *in vitro* model of zika infection (samples were cell culture media, Dulbecco’s modified Eagle’s Medium) and FAV contamination of river environment (river and sewage water samples were analysed) [21, 22].

The assay presented here, was developed and validated in accordance with Food and Drug Administration (FDA) guidelines, [23] with fundamental parameters assessed, including accuracy, precision and sensitivity. Criteria including linearity, accuracy (the degree of variation from known value, assessed by controls [QCs]), precision (the degree of variation within repeated measurements), selectivity (ensuring detection of the analyte and not an endogenous compound within the sample matrix), recovery (determining the percentage of recovery and more importantly the reproducibility of the extraction process) and stability were all assessed. The assay was primarily developed to investigate the pharmacokinetics of FAV in preclinical species (mouse plasma, phosphate buffered saline [PBS]). However, the assay was also validated for human plasma providing a much-needed tool for downstream pharmacokinetic-pharmacodynamic studies in clinical trials.

Despite the sensitivity and specificity of LC-MS/MS analysis, matrix effects are a well-documented source of major concern [24]. Matrix effects may impact various stages of the analytical process, such as ionisation of the analyte (either suppression or enhancement of ionisation) and extraction efficiency [24, 25]. Given the influence of the matrix on the quantification of an analyte, a change in matrix may have detrimental effects on the reliability of an assay. Therefore, the presented method was developed for robust quantification of FAV in multiple matrices. The majority of published methods describe quantification of FAV in a single matrix [26, 27]. The greatest advantage of the assay is robust quantification of FAV in multiple matrices, with minimal impact of matrix effect. The demonstrated versatility will allow assessment of FAV *in vitro* and i*n vivo.*

Given the global urgency to discover active therapeutics against SARS-CoV-2; and in light of the potential application of FAV, a robust LC-MS/MS method is described which enables the quantification of FAV in a range of biologically relevant matrices.

## Methods and Materials

### Materials

FAV and the internal standard (IS) emtricitabine (FTC) were purchased from Stratech Scientific Ltd (Cambridge, UK). Drug free mouse plasma and human plasma with lithium heparin were purchased from VWR International (PA, USA). LCMS grade methanol (MeOH) was purchased from Fisher Scientific (MA, USA). All other consumables were purchased from Merck Life Science UK LTD (Gillingham, UK) and were of LC-MS grade.

### Tuning for Favipiravir and Internal Standard

Detection of FAV was performed on a SCIEX 6500+ QTRAP (SCIEX MA, USA) operating in negative mode. FTC was selected as the IS due to similar log P (FAV 0.252, FTC −0.043) and had a similar retention time to FAV. Tuning was performed via direct infusion of FAV (10ng/mL at a flow rate of 10μL/min) to optimise compound-specific parameters (declustering potential, collision energy and collision exit potential) and source specific parameters (curtain gas, ionisation voltage, temperature, nebuliser gas and auxiliary gas).

### Chromatographic Separation

Samples were separated a multi-step gradient with a Kinetex® F5 column 2.1×100mm 2.6μm (Phenomenex CA,USA). Mobile phases A and B consisted of H_2_O with 0.1% acetic acid and MeOH with 0.1% acetic acid, respectively. A multi-step gradient was applied over 3 minutes at a flow rate of 600 μL/min as follows: initial conditions of 100% A and 0% B were increased over 2 minutes to 25% A and 75 % B. Mobile phase B was then increased to 99 % over 0.1 minutes and held for a further 0.5 mins. Gradient was restored to start conditions at 2.5 mins and help for 0.5 mins.

### Extraction from Mouse Plasma, human plasma and PBS

100 μl of standard, quality control (QC), blank or unknown sample was transferred to glass vials Samples were diluted with 500 μL of MeOH (containing 10ng/mL of IS) and thoroughly vortexed. Samples were then centrifuged at 3500g for 5 minutes at 4°C. 400 μL of supernatant fraction was transferred to a fresh glass vial and evaporated under a gentle stream of nitrogen. Samples were reconstituted in 100μl of H2O:MeOH (95:5). 50μl of the sample was then transferred into 200μl chromatography vials. 2μl of each sample was injected for analysis.

### Linearity

Calibrators were prepared by spiking untreated matrix with FAV followed by serial dilution, ranging from 0.78ng/mL to 200ng/mL. Linearity was assessed over 3 independent runs. Acceptance criteria were as follows deviation of standards interpolated concentration from stated concentration was set at 15%, excluding the lower limit of quantification (LLOQ) where deviation was set at no more than 20%. Acceptable R^2^ was set at >0.99.

### Recovery

To ensure the reproducibility of the extraction process, recovery was assessed at 3 quality control (QC) concentrations, 2ng/mL, 75 ng/mL and 150ng/mL independent of calibrators. Extracted samples were then compared to equivalent concentrations of un-extracted samples, representing 100%.

### Matrix Effects

The degree of interference from the matrix (due to potential interfering substances including endogenous matrix components, metabolites and decomposition products) was assessed by spiking extracted untreated matrix. Peak area was then compared to extracted LLOQ (0.78ng/mL). The LLOQ was a minimum of 5 times greater than the background signal. The assay was initially developed in mouse plasma and then further validated for mouse lung, PBS (used for bronchoalveolar) and human plasma.

### Accuracy and Precision

Accuracy and precision were assessed by preparation of three concentrations independent of the calibrators (2ng/mL, 75ng/mL and 150ng/mL). Deviation of mean values of each concentration must be within 15% of the stated concentration, with the exception of the lower concentration, where deviation must be less than 20%. Intra- and inter-assay variability of accuracy and precision were assessed to ensure robustness of the assay [25]. Full validation was assessed in mouse plasma. As a change of matrix is considered a minor change, the assay was part validated in human plasma and PBS assessing intra assay variability [25].

## Results

### Recovery of Favipiravir from Mouse Plasma

The extraction efficiency of FAV was determined at three QC concentrations using the optimised LC-MS/MS parameters (Table 1). The mean recovery was 76.5% (±1.76), 98.5% (±1.20) and 95.0% (±3.10) for mouse plasma, human plasma and PBS. Recovery varied between each matrix, however mean recovery across the 3 concentrations tested demonstrate the assay performance was highly reproducible (Figure 1).

**Table 1.** shows the optimised parameters used to detect FAV and FTC (IS). *denotes the daughter ion used for quantitation and ** denotes the daughter ion used for qualification.

| Parent (m/z) | Product (m/z) | RT (Min) | DP (Volts) | EP (Volts) | CE (Volts) | CXP (Volts) |
| --- | --- | --- | --- | --- | --- | --- |
| **Favipiravir** | *113.0 | 1.97 | 60 | 10 | 25 | 16 |
| 158.1 | **141.0 | 1.97 | 60 | 10 | 25 | 24 |
| **Emtricitabine (IS)** |  |  |  |  |  |  |
| 248.0 | 130.0 | 2.20 | 60 | 10 | 25 | 16 |
| Global Parameters | Curtain Gas | Collision Gas | Voltage (V) | Temperature (°C) | Gas 1 | Gas 2 |
|  | 25 | Medium | 4000 | 600 | 30 | 25 |

**Figure 1.**
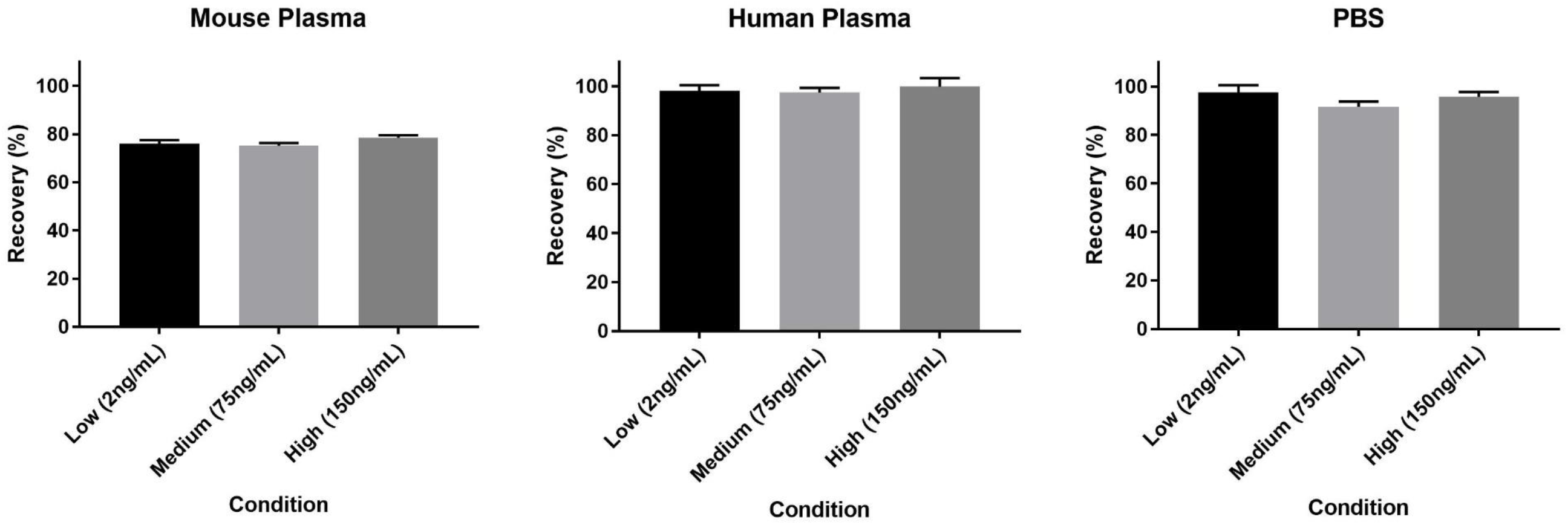
shows the percentage recovery for the low (a), medium (b) and high (c) QCs in extracted mouse plasma, human plasma and PBS. Data is show percentage of unextracted standards.

### Linearity

Extracted calibrators demonstrated strong linearity (mouse plasma R^2^ = 0.9989, human plasma R^2^ = 0.9971 and PBS R^2^ = 0.998) meeting all acceptance criteria (Figure 2). The calibration curve was fit to the data using a linear equation with sample weighting of 1/X.

**Figure 2.**
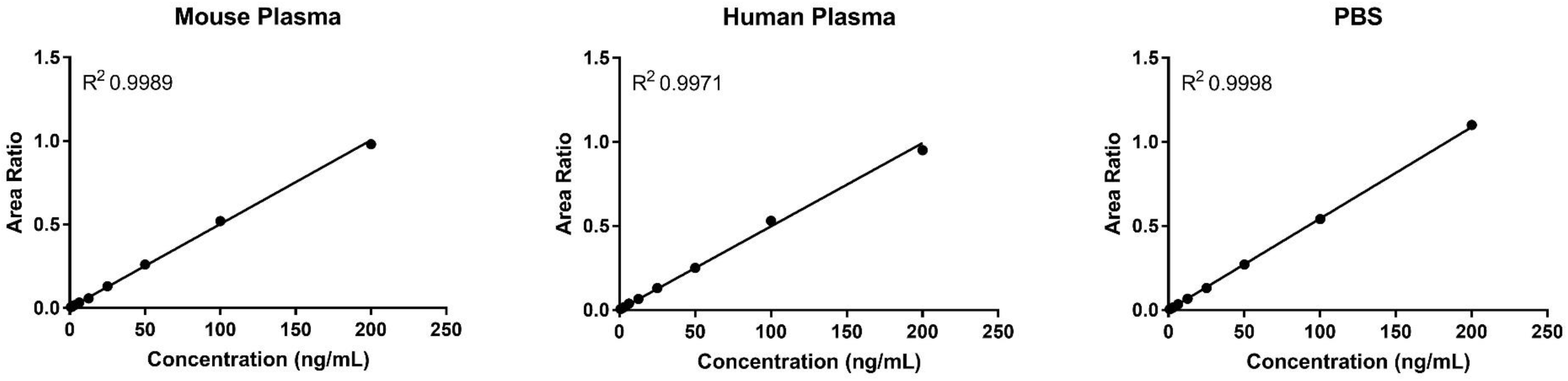
shows the standard curve generated from extracted mouse plasma, human plasma and PBS standards of FAV over the range of 0.78ng/mL to 200ng/ml.

### Selectivity

Matrix effects are well known to be a potential confounding factor in the development of LC-MS/MS assays. The matrix effect of mouse plasma, human plasma and PBS were determined using extracted blank matrix and compared to the peak area of the LLOQ (0.78ng/mL). The extracted blank showed no detectable peak at the retention time of FAV in mouse plasma, human plasma and PBS with a peak area of (Figure 3a, b and c). FDA guidelines require the signal produced by the LLOQ be greater than five-fold of the signal observed by the extracted blank (Figure 3d, e and F).

**Figure 3.**
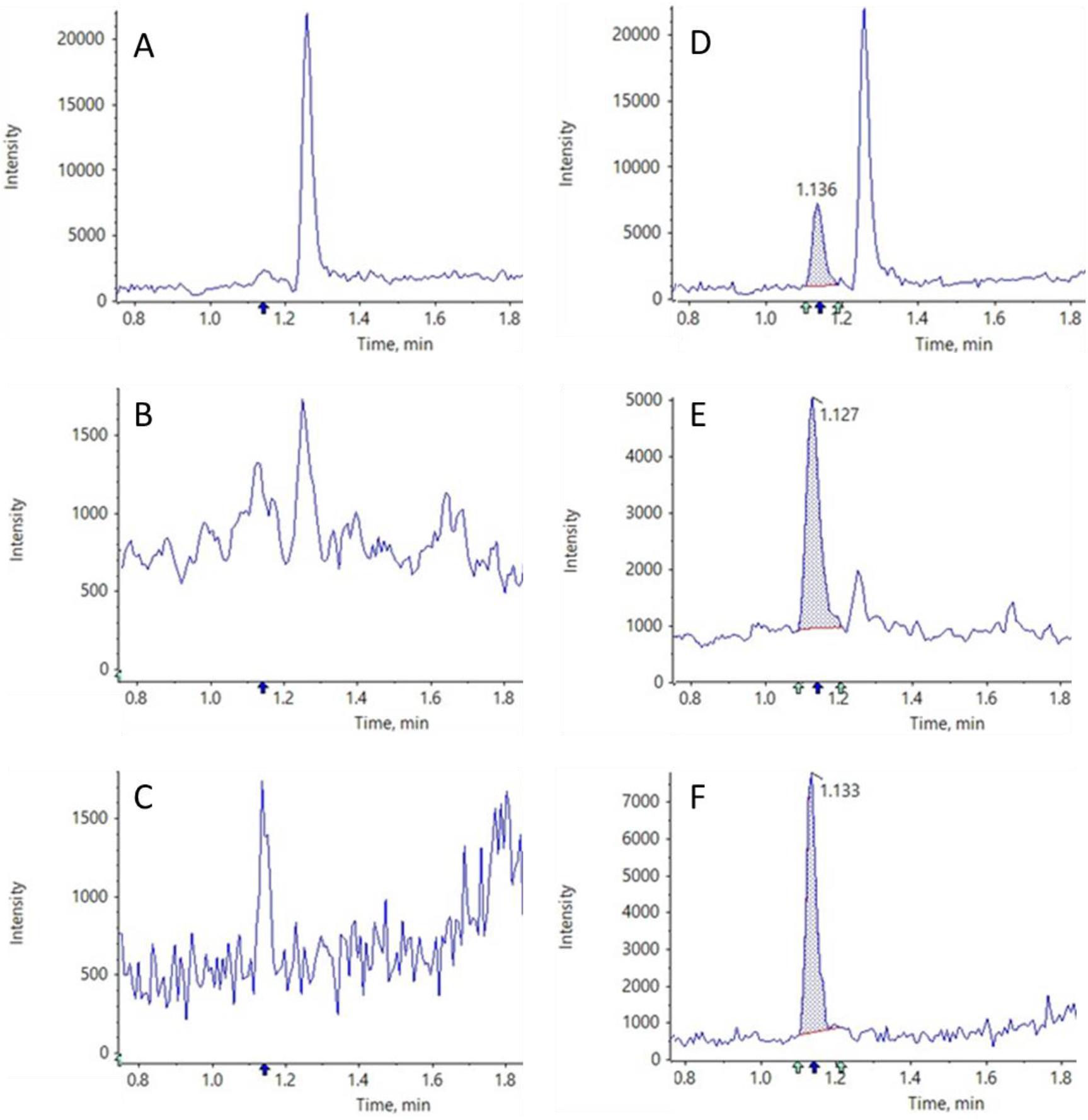
shows a representative chromatogram from blank mouse plasma (A), blank human plasma (B) and blank PBS (C). Also shown is a representative chromatogram of the LLOQ in mouse plasma (D), human plasma (E) and PBS (F). The retention time of FAV is also shown (1.1 min).

### Accuracy and Precision

The reproducibility and robustness of the assay was determined by examining the accuracy and precision of extracted QCs. Full validation was conducted using mouse plasma. The intra-assay % error in accuracy was below 15% at all levels, mean % error at 0.78ng/mL (0.13%), 75ng/mL (1.74%) and 150ng/mL (2.67%). The intra-assay percentage error of precision also fell below 15%, 0.78 ng/mL (6.16%), 75ng/mL (2.21%) and 150ng/mL (1.62%) (Table 2). The variability between assays was calculated to demonstrate that the assay-maintained accuracy and precision across repetitions of the assay. Table 2 shows inter-assay variance of accuracy and precision calculated from the mean values of the 3 repetitions of the assay. The percentage error in accuracy fell below 15% across all 3 repeats (range between 0.19% and 13.48%). Inter-assay percentage variance of precision also fell below 15% across all 3 repeats (range between 7.51% and 9.68%). In accordance with FDA guidelines human plasma and PBS required part validation [25]. The intra-assay % error in accuracy was below 15% at all levels in human plasma (mean % error at 0.78ng/mL 0.13%, 75ng/mL 1.74% and 150ng/mL2.67%) and PBS (mean % error at 0.78ng/mL 0.13%, 75ng/mL 1.74% and 150ng/mL2.67%). The intra-assay percentage error of precision also fell below 15% in human plasma (mean % error at 0.78ng/mL 0.13%, 75ng/mL 1.74% and 150ng/mL2.67%) and PBS (mean % error at 0.78ng/mL 0.13%, 75ng/mL 1.74% and 150ng/mL 2.67%, Table 3).

**Table 2.**
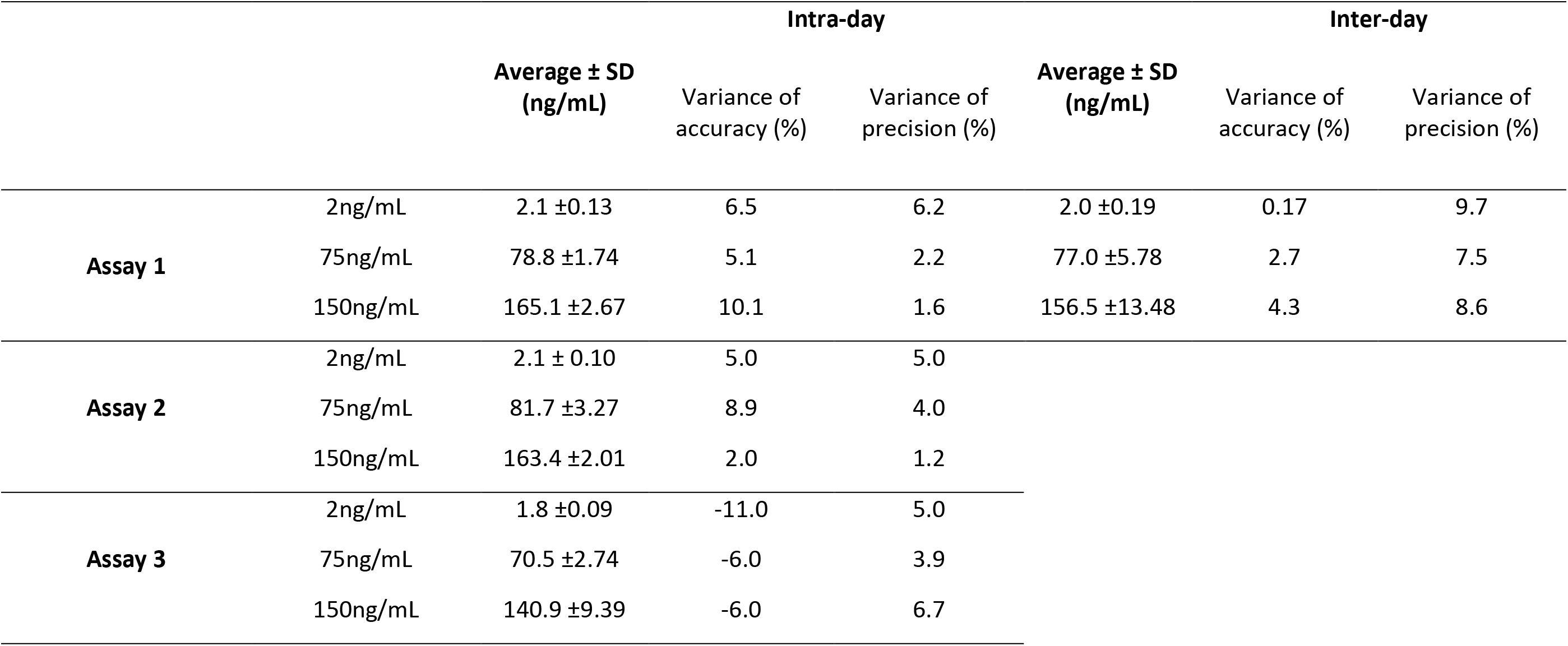
shows the intra-assay variance in accuracy and precision of 3 repetitions of the assay. Also shown is the variance in accuracy and precision of the inter-day assay performance. Accuracy and precision were assessed in triplicate at 3 levels (low (2ng/ml), medium (75ng/ml) and high (150ng/ml).

**Table 3.** shows the intra-assay variance in accuracy and precision of the assay in human plasma and PBS. Accuracy and precision were assessed in triplicate at 3 levels (low (2ng/ml), medium (75ng/ml) and high (150ng/ml).

|  |  | Intra-day |  |  |
| --- | --- | --- | --- | --- |
| | | Average ± SD<br>(ng/mL) | Variance of<br>accuracy (%) | Variance of<br>precision (%) |
| Human Plasma | 2ng/mL | 2.2 ±0.05 | 7.8 | 2.3 |
| | 75ng/mL | 77.6 ±1.79 | 3.5 | 2.3 |
| | 150ng/mL | 154.8 ±1.01 | 3.2 | 0.7 |
| PBS | 2ng/mL | 2.3 ±0.15 | 12.5 | 6.7 |
| | 75ng/mL | 77.1 ±3.08 | 2.8 | 4.0 |
| | 150ng/mL | 154.0 ±4.04 | 4.0 | 2.6 |

## Discussion

The current SARS-CoV-2 pandemic has led to an unprecedented global research effort to identify potential therapeutics for COVID-19 treatment or prevention. The repurposing of existing antiretrovirals that display opportunistic efficacy against SARS-CoV-2 presents opportunities for accelerated pharmacological interventions [12]. The identification of an antiretroviral already in use for another indication would dramatically cut the time required to identify and develop novel therapeutics. FAV was originally developed as an anti-influenza therapy but has been demonstrated to display efficacy against a range of viral infections, most recently being proposed for SARS-CoV-2 [11, 14]. The anti-SARS-CoV-2 activity of FAV requires more extensive evaluation *in vitro* and *in vivo,* but it has emerged as a potentially attractive candidate.

The presented work describes a rapid, robust and highly sensitive FAV LC-MS/MS assay for downstream applications. Quantification in mouse plasma demonstrated the assay to be highly reproducible, precise and accurate. Furthermore, the assay was able to quantify FAV from multiple diverse matrices, indicating a tolerance to the negative impact of matrix effects.

Initially, the assay was developed to quantify FAV in mouse plasma to support ongoing preclinical research. The final assay fully met the FDA bioanalytical method development guidelines, demonstrating good accuracy, precision and linearity. In addition to plasma, other biological compartments are of significant relevance to SARS-CoV-2. The assay was further validated in human plasma and PBS in order to facilitate clinical and *in vitro* investigations. As previously stated, matrix effects can have a detrimental impact on the performance of a LC-MS/MS assay. Coeluting endogenous compounds have the potential to negatively impact the ionisation of the analyte of interest, by either suppression or enhancement [28]. As FAV is 54% protein bound and slightly soluble in aqueous matrices (log P = 0.49), the change of matrix may lead to significant changes in the behaviour of the assay. In order to facilitate application of this assay by the wider research community, FAV was also quantified in human plasma. Despite the similarities between human and mouse plasma, species differences may lead to changes in the assay behaviour. Protein binding is commonly lower in preclinical species when compared to humans. Diazepam is 98% bound in human plasma and 84.8% bound in rats [29]. The species differences can be more profound, for example valproate is 94.8% bound in human but only 12% bound in mice almost 8 fold-difference [29]. The presented data demonstrates that matrix effects did affect the absolute recovery, but this did not affect the reproducibility and overall performance of the assay.

The presented assay is currently being adapted to quantify FAV from plasma from other preclinical species and other relevant *in vitro* matrices (cell culture media, foetal bovine serum and cell lysate). Additional work is also underway to adapt the assay for quantification in other tissues such as lung and nasal turbinate. However, due to the more complex nature of tissue homogenates considerable additional optimization may be required.

The presented assay surpasses many of the currently published methods for FAV detection. The most frequently used assays have been developed on HPLC with a LLOQ in the region of 100-1000ng/mL [17, 20]. Other methods have been developed for LC-MS/MS providing increased sensitivity in comparison to HPLC, with LLOQ in the region of 0.5-2000 ng/mL [21, 22]. In addition to greater sensitivity, the assay presented here demonstrates greater flexibility when applied to the analysis of a diverse array matrices relevant to the further exploration of FAV as a SARS-CoV-2 therapeutic.

In summary, optimisation of a robust, simple and sensitive LC-MS/MS assay for FAV is presented, which conforms to FDA bioanalytical development guidelines and was capable of assessing FAV in multiple preclinical and clinical matrices.

